# Help-seeking behavior for problematic substance uses in Bahir Dar town, North-West Ethiopia

**DOI:** 10.1101/363705

**Authors:** Habte Belete, Tesfa Mekonen, Wubalem Fekadu, Getasew Legas, Asmamaw Getnet

## Abstract

**Background:** Mental, neurological and substance use disorders are common, but 76% to 85% of people with those disorders in low and middle-income countries did not receive treatment.

**Objective:** Our objective was to assess the level of help seeking behavior and associated factors among residents with problematic substance uses (alcohol, khat, tobacco and hashish).

**Methods:** Community based cross sectional study was conducted in Bahir Dar town among total of 548 participants with problematic substance users. We had interviewed for help seeking behavior by pre-tested modified General Help Seeking Questionnaire. Logistic regression was done and p-value < 0.05 was used for declaration of significant level. Adjusted odds ratios (AOR) and 95% confidence intervals (CI) have been used.

**Results:** Among five hundred and forty-eight participants with problematic substance users, only one hundred and sixty-eight (30.7%) sought help for their substance related problems. Participants’ age above 35 years [AOR = .47 95% CI (.25, .90)], positively screened for common mental disorders [AOR = 4.12, 95% CI (2.7, 6.3)], comorbid medical [AOR = 3.0, 95% CI (1.7, 5.3)], and grand-families’ history of substance user [AOR = 2.18, 95% CI (1.4, 3.4)] found significantly associated with help seeking.

**Conclusion:** There was low proportion of help seeking behavior among participants with problematic substance users. Advanced age was a barrier to seek help while medical illnesses, common mental disorders and history of substance use in grand families were found to enforce to seek help.

## Introduction

Mental, neurological and substance use disorders are common; approximately one in four families has at least one member with a mental disorder which is accounted for 14% of the global burden of disease. Globally about 190 million drug users were reported and of them 40 million serious drug related illnesses or injuries were identified each year [1]. For example alcohol alone contributes for 7.6% males and 4% females of deaths whereas it contributes to more than 200 alcohol related diseases (alcohol related injuries, alcohol dependence, liver cirrhosis, and cancers) [2, 3]. However, 76% to 85% of people with Mental, neurological and substance use disorders in low and middle-income countries did not receive treatment for their behavioral problems [4-6].

Help-seeking is an adaptive coping process that is the attempt to obtain external assistance to deal with a mental health concern formally or informally. Formal help-seeking is assistance from professionals whereas informal help-seeking is assistance from informal social networks, such as friends and families [7]. In Ethiopia substance use in the community [8] and in university students has many social and academic related problems [9-11]. Despite the high prevalence of substance related problems, most people do not access professional health help. For instance, WHO reported that universally alcohol use disorder has a widest treatment gap (78.1%). However, problematic substance use is remaining a major public health concern and barriers to treatment for problematic substance uses have not been well searched in developing world [12]. There are evidences in which environmental factors serve as barriers to seek help for substance related behavioral disturbances in the community [13]. According to WHO report individual factors like, personal motivation, perception of need, lack of health insurance, internalized gender norms, and perceptions of social supports as positive, and others influence the help-seeking behaviour of adolescents’ [14]. From A community study among people with mental disorders including, alcohol use disorders, only 31.7% of them had sought help. From this report 15.7% help sought was from mental health providers, 8.4% from general practitioners, and 7.6% from religious/ spiritual advisors or other healers [15]. Professional help seeking behaviors among problematic drug user lesbian and bisexual women was only 41.5% [16]. Among people with alcohol use disorders 7% of them seek help from any mental health professional; 5.8% from medical health professional; 7.2% from a social support setting and 4.5% from any religious or spiritual advisor/healer [15]. Among patients who visited traditional healers for their mental health problems, 41% was visited to seek help for substance use disorders [17].

Factors related to enrolling in substance abuse treatment are multiple and researchers had blamed factors like lack of availability, transportation, and insurance [18]. Even though, men report higher levels of substance abuse than women and are more likely to have psychosocial problems, but are less likely to seek help [19]. Older adults’ positive attitudes and treatment beliefs were enabling to use mental health services [20]. Negative beliefs about the quality and effectiveness of treatment had a negatively effect to access services [21]. Reluctance to give up the substance; reluctance to admit the need for help and inability to afford treatment were contribute to poor help-seeking behavior [22]. Even if the magnitude of substance abuse in Ethiopia reaches 22.8% in the general population [23]; as per our best knowledge there is no publishing data to show prevalence and factors that associated with help seeking behaviour among problematic substance users in Ethiopia. Therefore, the purpose of this study was to see the level of help seeking behaviour and its contributing factor among problematic substance users in urban residents.

## Methods

### Study settings

We were conducted a community based cross-sectional study in Bahir Dar town which is located in Northwest Ethiopia around 565 kilo meters from Addis Ababa. The city has a total of 180,174 populations; of these 93,014 are females. Currently, there are four hospitals, ten health centers and a number of other private health institutions (clinics, pharmacies and drug shops). In this city alcohol and khat use are common among adolescents.

### Participants

Source population was all permanent residents in the city. Study population was those positively screened for problematic substance use and whose age 18 and above were including in the study. A participant who cannot communicate was not included in the study. Sample size was determined with a single population formula by using proportion of help seeking for problematic substance uses among residents 41% [17], 4% margin of error, and 95% confidence interval. We select four kebeles using lottery method from the total 17 kebeles. Then house-holds were selected proportionally from each selected kebeles. Finally the data has been collected at household level. Individual participants were selected based on their status of a problematic substance uses in the family. We interviewed 548 participants who were positive for problematic substance uses (Khat, alcohol, tobacco and cannabis) in the community. Data was collected by degree holder nurses with a self-administered questionnaire which was translated into Amharic version (local working language).

### Measurements

First the participants were screened for problematic substance uses by using CAGE AID questionnaire. CAGE AID has been developed to identify problematic substance uses in the community and has four items. Scoring of two or greater positive answers from the four questions to social drugs screening tool was considered as problematic substance uses [24]. CAGE-AID is derived from the four questions of the tool: Cut down, Annoyed, Guilty, and Eye-opener. Each has “yes and no” response that valued one point (1) and zero (0). This tool helps to screen the presence of problematic substance uses in the community. Help seeking was assessed if individuals sought assistance for their substance related problems from delineate formal and informal source [7]. Help seeking behavior among problematic substance users assessed by using modified General Help Seeking Questionnaire (GHSQ). It assesses whether help was sought or not and the potential sources of that help for the last 12 months. GHSQ has good validity, reliability, and reliability (chronbach’s alpha = 0.83) [25] and we did a pre-test for our study and has internal consistency (chronbach’s alpha = 0.80). Social support assessed by using oslo-3 social support scale that consist three items [26]. Common mental disorder was considered when adults who score 11 or more symptoms of the 20 self-reporting questionnaire in the last one month. The self-reporting questionnaire developed to screening the presence of common mental disorders among community participants and developed by WHO [27]. Income was assessed by using relative income by leveling their own income as less than others; similar to others and better than others. Education was categorized in uneducated (who were unable to read and write); and educated (which were able to read and write). Clinical variables like, co morbid diagnosis medical illnesses were assessed by asking the participants, if they had a diagnosis of medical illnesses before the survey.

### Analysis

After the data was checked for completeness and consistency, it was coded and entered in the Epi-Data 3.1 software. Data was exported to SPSS for analysis and p-value less than 0.05 was declaration of a statistically significant. Bivariate and multivariate logistic regression analyses had done to identify determinants of help seeking behavior for substance use related problems.

### Ethical clearance

Ethical clearance was obtained from Ethical Review committee of college of medicine and health sciences, Bahir Dar University. Formal permission letter was taken from administrative of the city and written consent was taken from the participants. All participants who were problematic substance users referred to clinic for better screening to mental and medical illnesses. Those who were positive for screening of common mental disorders were link with psychiatric clinics

## Results

A total of 548 residents who problematic substance uses were included in the study and 422 (77%) of them were males. The median age of participants was 27 years and substantial number of them were living alone, 241 (44%) (table1).

**Table1:**
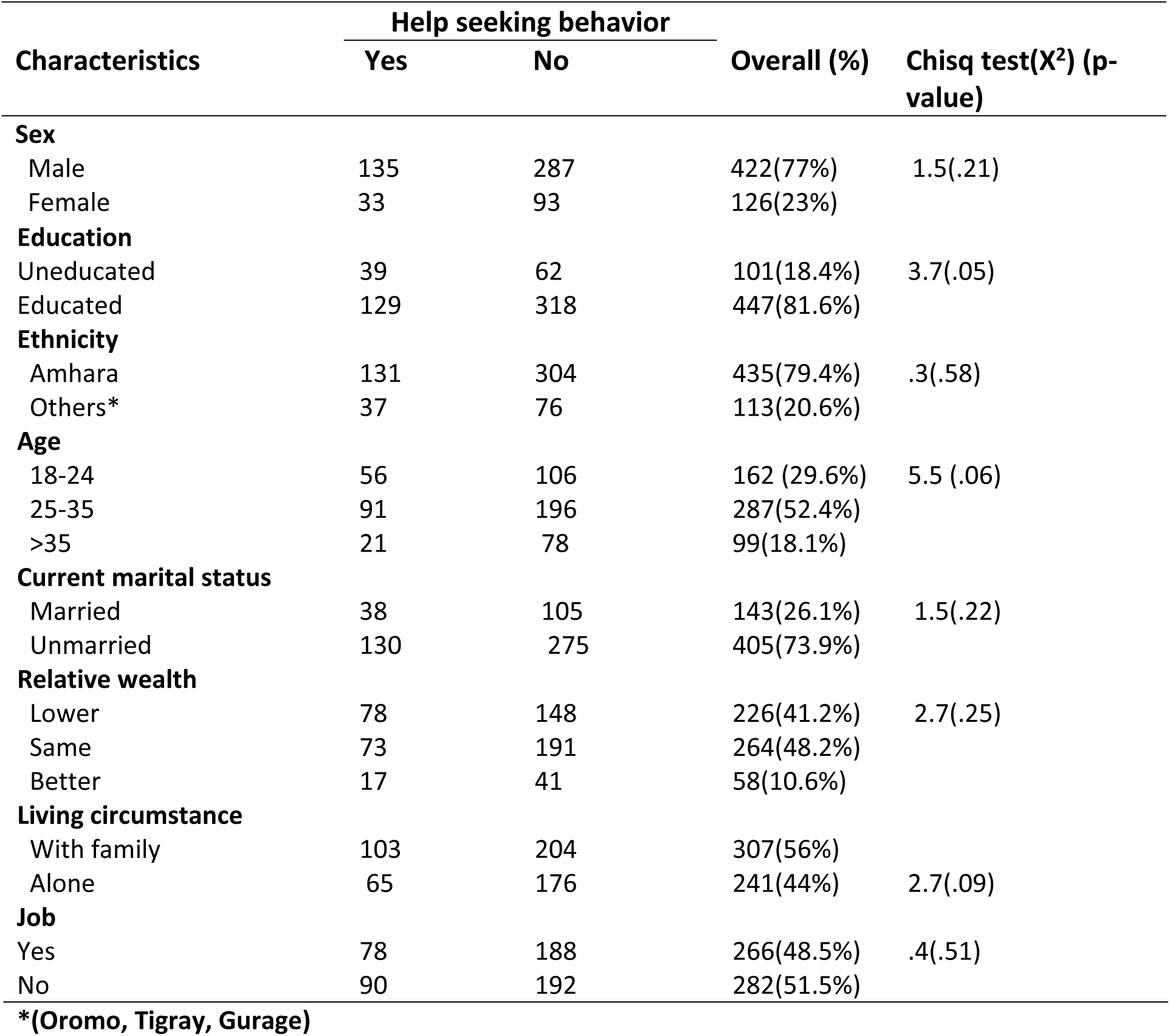
Socio-demographic characteristics and help seeking behavior among problematic substance users in North-West Ethiopia (n=548)

From total participants one hundred sixty one (29.4%) had substance user family members; two hundred forty (37.2%) had substance user friends and three hundred nineteen (58.2%) had history of substance use among their grandparents. Two hundred three (37%) problematic substance users had found with poor social support and one hundred forty nine (27.2%) had common mental disorders (table 2).

**Table 2:** psychosocial factors related to help seeking behavior among problematic substance users in North-West Ethiopia (n=548)

| Characteristics | Help seeking behavior |  | Overall (%) | Chisq test (X<sup>2</sup>) (p-value) |
| --- | --- | --- | --- | --- |
|  | Yes | No |  |  |
| History of substance use in grand family |  |  |  |  |
| Yes | 125 | 194 | 319(41.8%) | 26.1(.000) |
| No | 43 | 186 | 229(58.2%) |  |
| Who had substance user friend |  |  |  |  |
| Yes | 78 | 126 | 204(37.2%) | 8.8(.003) |
| No | 90 | 254 | 344(62.8%) |  |
| **Substance use leads economic crisis** |  |  |  |  |
| Yes | 124 | 269 | 393(71.7%) | .5(.46) |
| No | 44 | 111 | 155(28.3%) |  |
| **Common mental disorders** |  |  |  |  |
| Yes | 85 | 64 | 149(27.2%) | 67.0(.000) |
| No | 83 | 316 | 399(72.8%) |  |
| **Medical illnesses** |  |  |  |  |
| Yes | 38 | 33 | 71(13%) | 20.0(.000) |
| No | 130 | 347 | 477(87%) |  |
| **Think on substance cause mental illness** |  |  |  |  |
| Yes | 131 | 278 | 409(74.6%) | 1.4(.23) |
| No | 37 | 102 | 139(25.4%) |  |
| **Level of social support** |  |  |  |  |
| Poor | 71 | 132 | 203(37%) | 3.8(.15) |
| Moderate | 81 | 195 | 276(50.4%) |  |
| Good | 16 | 53 | 69(12.6%) |  |

### Magnitude of help seeking behavior

Only one hundred and sixty eight (30.7%) sought help for their substance related behavioral disturbance among five hundred forty eight problematic substance users. Most of them were sought help from the informal source. Among these one hundred and twenty four (22.6%) sought help from their love; one hundred ten (20.1%) from their friends; one hundred and three (18.8%) from their families; eighty four (15.3%) from their relatives; and one hundred (18.2%) from religious institutes. From the formal help seekers ninety (16.4%) sought help from mental health professionals; and eighty nine (16.2%) from general medical practitioner.

In terms on their mental health status among help seekers, 50.6% had common mental disorders, 42.3% had poor social support and 9.5% had strong social support. Among help seekers gender difference was examined and 32% male and 26.2% females were sought help.

### Multivariate analysis

After bivariate and multivariate analysis of help seeking behavior in relation to all independent variables; advanced age, comorbid medical illnesses, had grand-parents with history of substance use, and positively screened for common mental disorders were found to be statistically significant (table 3).

**Table 3:** Factors association with help seeking behavior among problematic substance users, North-West Ethiopia (n=548)

| Variables | Help seeking behavior |  | (Crude odd ratio)<br>(95% CI) | AOR<br>95% CI | p-value |
| --- | --- | --- | --- | --- | --- |
|  | Yes | No |  |  |  |
| **Age in years** |  |  |  |  |  |
| 18-24 | 56 | 106 | 1.00 | 1.00 |  |
| 25-35 | 91 | 196 | .88 (.58, 1.32) | .89 (.57, 1.41) | .63 |
| >35 | 21 | 78 | .51 (.28, .91) | .47 (.25, .90)* | .022 |
| **Common mental disorders** |  |  |  |  |  |
| Yes | 85 | 64 | 5.0 (3.37, 7.58) | 4.12 (2.7, 6.3)* | .000 |
| No | 83 | 316 | 1.00 | 1.00 |  |
| **Medical illnesses** |  |  |  |  |  |
| Yes | 38 | 33 | 3.1 (1.85, 5.11) | 3.00 (1.69, 5.34)* | .000 |
| No | 130 | 347 | 1.00 | 1.00 |  |
| **History of substance use in grand family** |  |  |  |  |  |
| Yes | 125 | 194 | 2.8 (1.87, 4.16) | 2.18 (1.42, 3.37)* | .000 |
| No | 43 | 186 | 1.00 | 1.00 |  |
| **Had substance user friend** |  |  |  |  |  |
| Yes | 78 | 126 | 1.7 (1.2, 2.5) | 1.43(.95, 2.16) | .087 |
| No | 90 | 254 | 1.00 | 1.00 |  |

\* (significance at P value <.05) and 1.00 (reference)

## Discussion

Even though 190 million drug users were reported globally and 40 million of them suffer from serious drug related illnesses each year [1], we found only 30.7% of the participants were sought help for their substance related behavioral disturbance. The care of people with behavioral disorders is a growing public health concern in the globe. However, 76% to 85% of people with mental, neurological and substance use disorders in low and middle-income countries not receive treatment [4-6]. This study coincides with the previous global report which stated low treatment seeking behavior in poor income countries. This result coincides with study done in Singapore 31.7% [15], but less than Los Angeles study 41.5% [16] and South African study 41% [17]. Even formal help seeking behavior in this study (16.4% from mental health professionals) found similar with Singapore study (15.7% from mental health providers) [15]. However, help seeking from general medical practitioner found higher in this study (16.2%) than Singapore (8.4%). Even though many psycho-social determinants accountable for the discrepancy of magnitude of help seeking behaviour, generally modern help seeking behavior looks low in developing countries.

Participants’ age above 35 years found negatively associated with help seeking behavior and decreased by 53% of their need to get help [AOR = .47 95% CI (.25, .90)]. This contradicted with a previous study that stated older adults were seeking more mental health services than Youngers [20]. Elder adults may not have a good family support to motivate them to get assistance for their behavioral disturbance. Participants who screened positively for common mental disorders found four times more help seeker than those who hadn’t common mental disorders [AOR = 4.12, 95% CI (2.7, 6.3)]. This may be due to the co-morbid effect of common mental disorders with substance related behavioral disturbance on their daily activities that leads to unable to control by their own capacity. The dual effect on their mental and behavioral status become severs and may push them to seek help. This finding agrees with the studies done in America [28, 29]. Comorbid diagnosis medical illness was identified as the determinant factor for help seeking behavior among problematic substance users.

Problematic substance users with comorbid medical illnesses were three times more likely to sought help than participants free of comorbid medical illnesses [AOR = 3.0, 95% CI (1.7, 5.3)]. This comorbidity may result perceived fear of death or other complications among participants and/or family members which give alarm to sought help from others. This finding agrees with previous study in America [30].

History of substance use in grand-families had a significant association with help seeking behavior among problematic substance users. Help seeking behavior found two times more common among problematic substance users who had grand-families with history of substance use than who hadn’t [AOR = 2.18, 95% CI (1.4, 3.4)]. The historical impact of substance related behavioral disturbance in the family system (in terms of morbidity and mortality) may perceived by the family members and this might motivated substance users to sought help to prevent further disabilities in the family.

### Conclusion and policy implication

This finding comes with low proportion of help seeking behavior among problematic substance uses for their substance related behavioral disturbances. It was found in line with WHO global report which stated 76% to 85% of people with Mental, neurological and substance use disorders in low and middle-income countries did not receive treatment. Advanced age was found a barrier to seek help while common mental disorders and history of substance use in grand families were found enforce to seek help. It is better to create awareness for potential complications of problematic substance uses and encourage substance users to visit mental health service in the community.

### Strengths and limitations of this study

We had tried to select participants with problematic substance users that non-problematic substance users. We have a limitation and the study was conducted in urban and lack of representation for the rural settings which may be the least help seeking behaviour can observed. We recommended for future studies shall be conducted in the rural part of Ethiopia.

## Acronyms

CAGE: Cut down, Annoyed, Guilty, and Eye-opener
GHSQ: General Help Seeking Questionnaire
SPSS: Stasticaly Package for Social Science
USA: United State of America
WHO: World Health Organization

## Ethics approval and consent to participate

Ethical clearance was obtained from Ethical Review Committee of college of medicine and health sciences, Bahir Dar University. Written consent taken from the participants.

## Consent for publication

Not applicable

## Availability of data and material

Data sets are available with author for reasonable request

## Authors’ contributions

All authors perform the design of the study, statistical analyses and draft the manuscript. All authors review and approve the final manuscript

## Acknowledgements

We acknowledge the study participants for their genuine participations

